# Navigating infection risk during oviposition and larval foraging in a holometabolous insect

**DOI:** 10.1101/171132

**Authors:** Jonathon A. Siva-Jothy, Katy M. Monteith, Pedro F. Vale

## Abstract

Deciding where to eat and raise offspring carries important fitness consequences for all animals, especially if foraging, feeding and reproduction increase the risk of exposure to pathogens. In insects with complete metamorphosis, foraging occurs mainly during the larval stage, while oviposition decisions are taken by adult-stage females. Selection for infection avoidance behaviours may therefore be developmentally uncoupled. Using a combination of experimental infections and behavioural choice assays, here we tested if *Drosophila melanogaster* fruit flies avoid potentially infectious environments at distinct developmental stages. When given conspecific fly carcasses as a food source, larval-stage flies did not discriminate between carcasses that were clean or infected with the pathogenic Drosophila C Virus (DCV), even though scavenging was a viable route of DCV transmission. Adult females however, discriminated between different oviposition sites, laying more eggs near a clean rather than an infectious carcass if they were healthy; DCV-infected females did not discriminate between the two environments. While potentially risky, laying eggs near potentially infectious carcasses was always preferred to sites containing only fly medium. Our findings suggest that infection avoidance can play an important role in how mothers provision their offspring, and underline the need to consider infection avoidance behaviours at multiple life-stages.

## Introduction

The behavioural immune system, the suite of behaviours that allow animals to avoid contact with infectious environments or conspecifics, is the first line of defence against infection [1–3]. Avoidance of infection relies on detecting cues of parasite presence - such as visual cues of infection risk or secondary pathogen metabolites – and integrating this sensory information to avoid sources of infection [4–10]. In addition to external cues of infection risk, the internal state of the animal, including its physiological status as a result of prior pathogen exposure, may also affect the ability to detect and avoid infection [11–13].

Avoiding contact with pathogens allows healthy individuals to escape the pathology that results from infection, and also prevents the deployment of the immune response, which may be metabolically costly and even cause immunopathology[2,3,14]. Despite these clear advantages, avoiding infection completely is rarely possible. Foraging and feeding, for example, are vital aspects of host metabolism, and are key to organismal reproduction and fitness, but they are also major routes of pathogen transmission [15,16].

Foraging and feeding are particularly important for holometabolous insect larvae, which devote most of their time to these behaviours. In situations of severe nutritional scarcity, larvae may even resort to cannibalism. For example, larvae of the fruit fly *Drosophila melanogaster* readily eat the carcasses of conspecifics following periods of starvation [17,18]. Cannibalism may appear to be a beneficial strategy when the alternative is starvation, but may increase the risk of trophic transmission of pathogens and parasites, especially if infected individuals are more likely to be targeted for cannibalism. While larvae of many insect species are frequently observed to avoid infectious environments or food sources [19], it is currently unclear if trophic infection avoidance occurs during cannibalistic scavenging.

Beyond foraging during the larval stage, choosing where to oviposit or rear offspring is another important life-history decision, but can be risky if individuals are unable to identify and avoid potentially infectious environments. The environment in which adult insects choose to oviposit is therefore a major determinant both of offspring environmental quality and infection risk [7,16,20]. Infection avoidance by insects during oviposition has been observed in response to a number of parasites and appears to be driven by diverse sensory cues, including avoidance of parasitoid wasp visual cues [7], and olfactory detection of bacteria and fungi [6,10]. Together, both adult oviposition choice and larval food preference determine the likelihood of infection in the early life-stages of holometabolous insects, and therefore both behaviours play an important role in disease transmission dynamics [4,21].

Here, we investigate larval foraging and adult oviposition in a holometabolous insect - the fruit fly *Drosophila melanogaster* - in the context of infection avoidance. Our study consisted of choice assays performed on either larval or adult stage *D. melanogaster*. Fly larvae were presented with a choice of scavenging on either a clean, non-infectious adult fly carcass, or a carcass that had been previously inoculated with a systemic Drosophila C Virus (DCV) infection (Figure 1a). In a second experiment, we tested adult oviposition choice by giving female flies the choice to lay eggs on a clean food source, a clean food source also containing a clean carcass, and a food source containing a carcass with a systemic DCV infection (Figure 1b). This 3-way choice assay allowed us to examine an important conflict faced by mothers: a carcass may present an additional nutritional source for future offspring, but may also present a potential risk of infection. In both experiments we assessed the fitness consequences of choices at both life-stages by following the development and longevity of larva (or laid eggs) as adult flies.

**Figure 1 - Experimental design.**
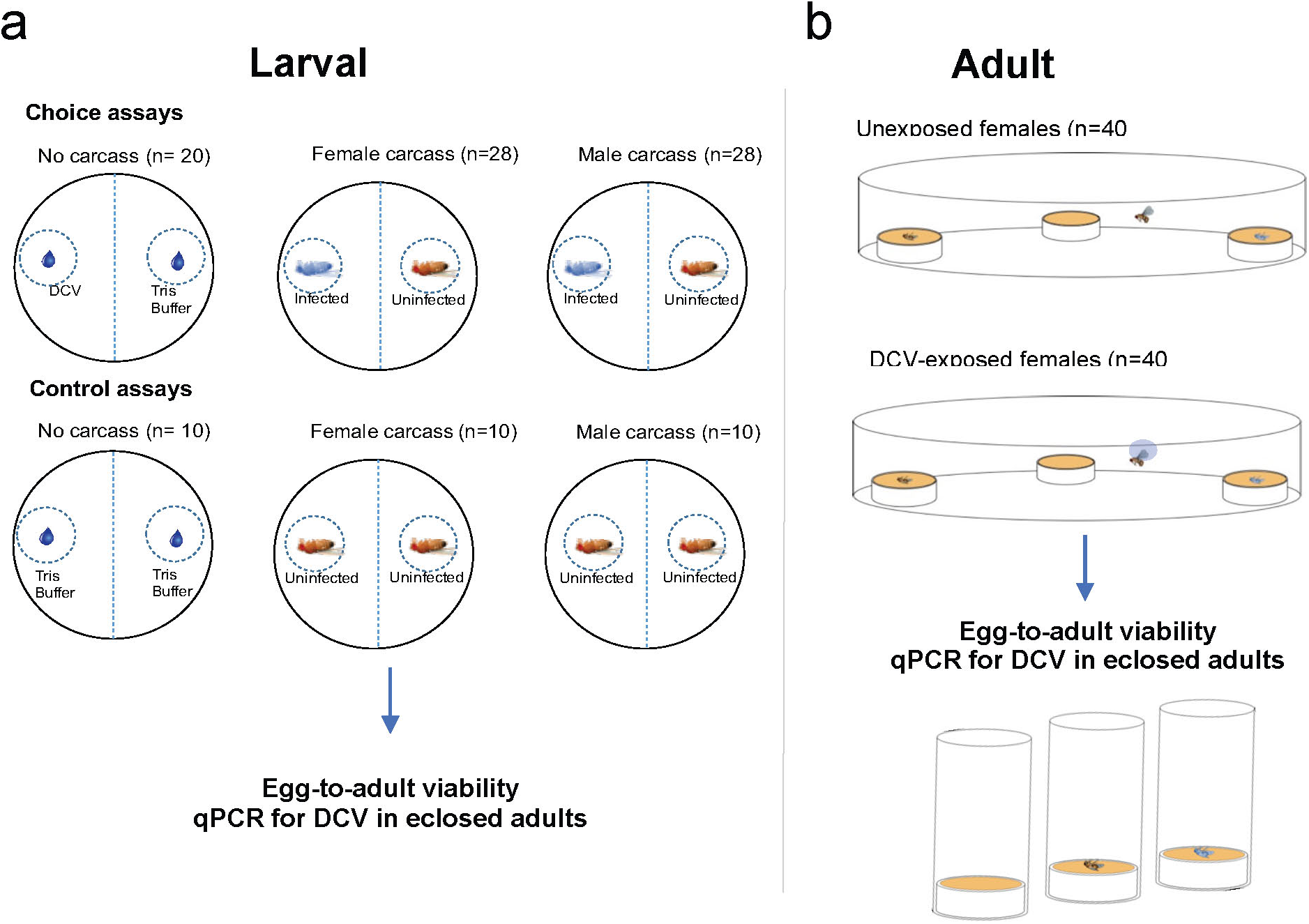
(a) Two-choice chamber used to measure larval foraging choice when presented with infectious and non-infectious food sources and the life-history data collected after the 72-hour assay. Petri dishes were set up as either two-choice plates (containing an infectious and non-infectious food source) or control plates (containing only non-infectious food sources). Eggs were placed at the centre of each plate, allowed to hatch and left for 72 hours whereupon the position of larvae was recorded to assay infection avoidance. (b) Three-choice chamber used to assay oviposition site choice in infected and uninfected mothers when presented with three sites containing just food, food and a fly carcass and food and an infected fly carcass. The number of eggs laid at each site was measured twice at two 24 hour intervals. After 48 hours, oviposition sites were removed and clutches were allowed to develop to adults whereupon the viral load of a randomly selected sub-sample was assayed.

## Materials and Methods

### Fly lines and rearing conditions

Both experiments used laboratory stocks of *D.melanogaster* Oregon R (OreR). Fly stocks were kept in plastic bottles (6oz; Genesee Scientific, San Diego, California, US) on a standard diet of Lewis medium [22] at 18±1˚C with a 12 hour light:dark cycle. Stocks were tipped approximately every 21 days into new bottles. Before the experiments, flies were transferred to clean bottles and maintained at low density (~50 flies per bottle) for a minimum of two generations at 25±1˚C with a 12 hour light:dark cycle.

### Virus culture and infection

*Drosophila* C Virus (DCV) is a horizontally transmitted positive-sense ssRNA virus of the Dicistroviridae family [23]. DCV infection establishes in the digestive, reproductive and fat tissues, resulting in a range of behavioural and physiological pathologies in both larval and adult stage flies, including reduced locomotor activity, metabolic and reproductive dysfunction, and eventually death [24–28]. The DCV isolate used in this experiment was originally isolated in Charolles, France [29] and was grown in Schneider Drosophila Line 2 (DL2) as previously described [27], serially diluted ten-fold in TRIS-HCl solution (pH=7.3), aliquoted and frozen at −80˚C until required. To infect flies, Austerlitz insect pins (0.15mm in diameter) were bent at a 90º angle ~0.5mm from the tip, dipped in DCV (10^8^ infectious units (IU) per ml), and inserted into the pleural suture of flies under CO₂ anaesthesia. Control infections employed the same protocol but with a needle tip dipped in sterile TRIS solution.

### Infection avoidance during larval foraging

We first tested if fly larvae could discriminate between healthy and potentially infectious fly carcasses. To generate these carcasses 4-7 day old male and female flies were randomly selected from an age-matched population. For each sex, half of the flies were stabbed with DCV 10⁷ DCV copies/ml and the other half stabbed with sterile TRIS buffer. Following 6 days (to allow viral replication), flies were frozen at −80 ˚C until required. We confirmed the infection status of the carcasses using DCV-specific qRT-PCR(see below) by randomly picking 5 male and 5 female flies.

We carried out a two-choice assay by placing ~100 fly eggs at the centre of each Petri dish containing ~20ml solid agar (5% sugar), and allowed the resulting 3^rd^ instar larvae to forage towards either a clean fly carcass or a carcass infected with DCV, placed at an equidistant positon from the eggs (3cm). We set up 56 ‘choice’ assays where larvae could choose between a clean or DCV infected carcass, and 20 ‘control’ assays, where both carcasses were clean (half of assays contained male carcasses, and the other half contained female carcasses). To differentiate between any effects of carcass degradation from a direct effect of DCV presence on larval choice, we also set up an additional 30 plates without fly carcasses, containing 10*µ*l of DCV (10⁷DCV IU/ml) and 10*µ*l of TRIS (two-choice; N=20) or only TRIS (control; N=10). 18 of the 106 plates set up across all treatments were excluded from the final dataset due to damage to the agar discriminating larval movement and thus providing unreliable results. All assays were conducted at 25±1˚C with a 12-hour light:dark cycle before being photographed after 72 hours. Images were marked using Adobe Photoshop CS3 to count the number of larvae within each plate half and within an area immediately surrounding the carcasses/droplets (~2.2cm in diameter – see Figure 1a).

### Larval infection status and virus quantification

We randomly selected 10 wandering-stage larvae found immediately adjacent to each carcass in 20 ‘choice plates’ and one carcass in 6 ‘control plates’ to assess DCV infection status and quantify viral load. Viral quantification was carried out by absolute quantification of DCV RNA copies using qRT-PCR. Total RNA was extracted by homogenising the flies or larvae in TRI Reagent (Invitrogen, Carlsbad, California, US) and using Direct-zol RNA miniprep kit (Zymo Research, Irvine, California, US), including a DNase step. The eluted RNA was then reverse-transcribed with M-MLV reverse transcriptase (Promega, Madison, Wisconsin, US) and random hexamer primers, and then diluted 1:1 with nuclease free water. The qRT-PCR was performed on an Applied Biosystems StepOnePlus system using Fast SYBR Green Master Mix (Applied Biosystems, Foster City, California, US) using the following forward and reverse primers, which include 5’-AT rich flaps to improve fluorescence [30] (DCV_Forward: 5’ AATAAATCATAAGCCACTGTGATTGATACAACAGAC 3’; DCV_Reverse: 5’ AATAAATCATAAGAAGCACGATACTTCTTCCAAACC 3’; with the following PCR cycle: 95°C for 2min followed by 40 cycles of: 95°C for 10 sec followed by 60°C for 30 sec. Two qRT-PCR reactions (technical replicates) were carried out per sample. For absolute quantification of DCV, the concentrations of DCV in the samples were extrapolated from a standard curve created from a 10-fold serial dilution (1-10^-^6) of DCV cDNA.

### Larval development and infection status

To analyse the effect of foraging choice on larval development, we removed 15 larvae found within 2cm of each carcass from 20 ‘choice’ plates and from one carcass on 6 ‘control’ plates. Larvae from each carcass were transferred together into plastic vials containing Lewis medium and we recorded the number of larvae that developed into pupae and the number of eclosed adults.

### Infection avoidance during oviposition

Following our test of infection avoidance at the larval stage, we tested the oviposition preference of female *D. melanogaster* when presented with a choice of clean and potentially infectious oviposition sites. Choice chambers were constructed by joining two lids of transparent plastic Petri dishes with adhesive tape, making a chamber 10cm in diameter and 2 cm in height. Chambers contained three oviposition sites comprised of upturned caps filled with Lewis medium, arranged in a triangle, each site, 50mm from the other two (Figure 1b). Oviposition sites contained either only Lewis medium, Lewis medium and an uninfected fly carcass, or Lewis medium and a DCV-infected fly carcass (infection protocol described above).

Three-day-old flies (*N*=40 males and 40 females) were isolated as virgins and stabbed with a virus-contaminated or sterile, virus-free control solution. Following infection, flies to be used in the oviposition assay were introduced to two males for mating for 72 hours. We then introduced a single mated female fly to each chamber and placed at 25°C (12-hour light:dark cycle) to allow oviposition. Two females (1 infected and 1 uninfected) laid no eggs during the experiment so were excluded from the final dataset. In total, we analysed the oviposition choice of 78 females. As DCV has been reported to affect *D. melanogaster* fecundity, to account for differences in the total number of eggs laid by our infection treatment group we measured oviposition site choice by counting the number of eggs at each site rather than the proportion of eggs laid at the three respective sites. To count the number of eggs laid on each oviposition site, photos were taken of individual oviposition sites with a Leica MC170 HD camera attachment on a Leica 0.32x/WD 200mm S8APO microscope (Leica microsystems, Wetzlar, Germany) after 24 and 48 hours.

### Fitness consequences of oviposition site choice

We quantified the potential fitness consequences of oviposition preference by transferring all oviposition sites, including carcass (if present), to individual vials and recorded egg-to-adult viability. Adults that eclosed from clutches during this experiment were frozen alongside in TRI reagent and DCV infection analysed using the same protocol as above. A total of 24 clutches were analysed in this way, with 6 oviposition sites excluded due to degradation or contamination during qPCR preparation.

### Statistical Analyses

In the larval choice experiment, we analysed the proportion of larvae choosing a given plate half or carcass area; larval DCV titers; the proportion of larvae developing into pupae (logit transformed); and the proportion of pupae that developed into adult flies (logit transformed). All response variables were analysed using Generalised Linear Models (GLMs) with ‘carcass sex’ and ‘carcass infection status’ and their interactions as fixed effects. In the adult oviposition experiment, we used the number of eggs laid at each oviposition site to assess infection avoidance. We analysed eggs counts, rather than the proportion of eggs laid on each oviposition site, to account for potential differences in fecundity between infected and uninfected flies (e.g. [28,31]. The number of eggs laid in the two measuring periods (0-24 hours and 24-48 hours) was analysed separately using generalised linear mixed models (GLMM) with Poisson distributed error. The oviposition site, infection status of the fly as well as an interaction between the two were listed as fixed effects. The total number of eggs laid and the choice chamber were included as random effects, with the latter nested within the fly’s infection status, to account for repeated measures and non-independence. The proportion of eggs that later eclosed as adults (egg-to-adult viability) was analysed using a GLMM with a binomially distributed error, with oviposition site included as a fixed effect. All statistical analyses and graphics were carried out and produced in R 3.3.0 using the *ggplot2*, *lme4* and *multcomp* packages.

## Results

### Larval flies do not avoid infectious food sources when scavenging

Fly larvae that hatched from eggs placed in the centre of the Petri dish, dispersed towards and consumed the fly carcasses placed at the edges of the dish (**Video S1**). We found no evidence that fly larvae can avoid infected food sources. Regardless of the measure of preference (plate half larvae were found in or the area surrounding each carcass or TRIS droplet) larvae showed no significant preference for clean or infected fly carcasses (Figures 2a, 2b; Table 1).

**Figure 2 – Larval foraging choice.**
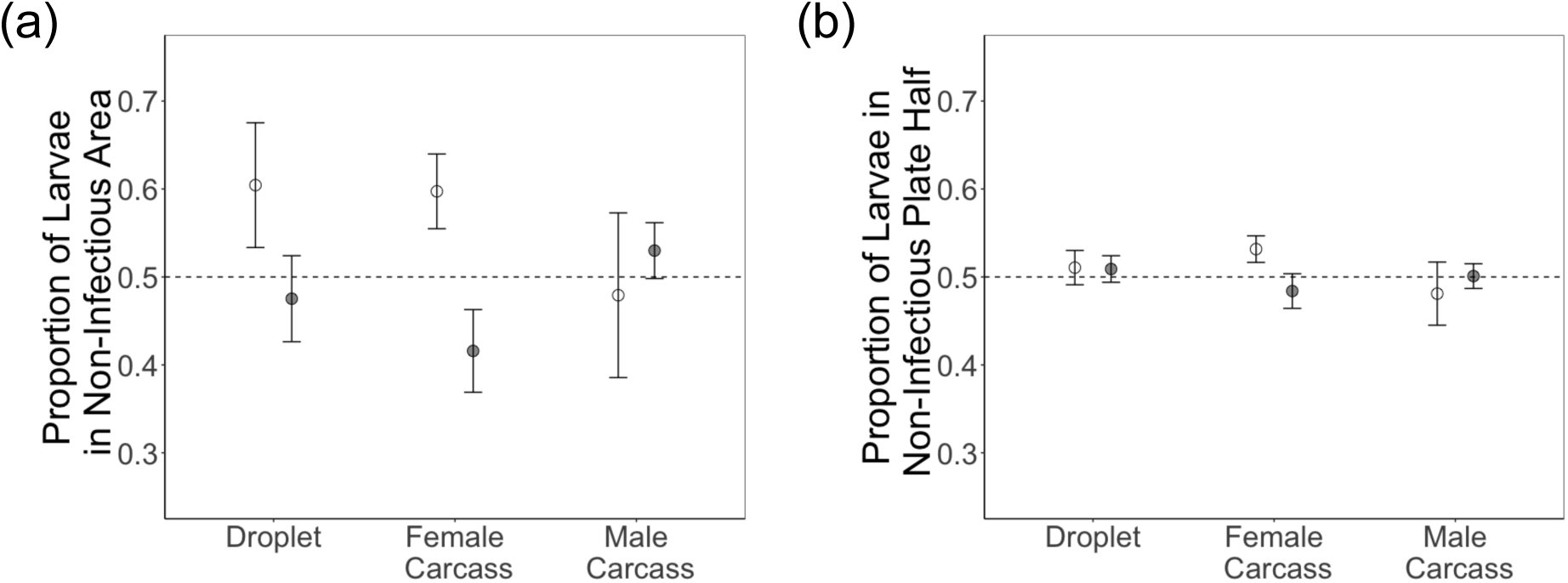
Mean±SE proportion of larvae on choice plates after 72 hours found (a) within area 2.2cm in diameter of the non-infectious food source and (b) on the non-infectious food source’s half of the plate. Results from both choice (white points) and control plates (grey points) are shown. In the case of choice plates, where only non-infectious food sources are present, the mean±SE is derived from the proportion of larvae present at a randomly selected side of the plate. Food sources included droplets of TRIS, a male carcass or female carcass.

**Table 1.** Model outputs for statistical tests performed on all experiments testing the causes and costs of infection avoidance in D. *melanogaster* larval foraging. Significant predictors are marked with asterisks (p<0.05=*, p<0.01=** and p<0.001=***).

| Response Variable | Predictor | DF | $\chi^2$ | p-value |
| --- | --- | --- | --- | --- |
| **Larval Foraging Choice by Plate Half** | Carcass Sex/TRIS | 2 | 0.5991335 | 0.7411 |
|  | Carcass Infection Status | 1 | 0.6326616 | 0.4264 |
|  | Carcass Sex/TRIS * Carcass Infection Status | 2 | 2.7615792 | 0.2514 |
| **Larval Foraging Choice by Carcass Area** | Carcass Sex/TRIS | 2 | 0.5124854 | 0.7740 |
|  | Carcass Infection Status | 1 | 3.5960234 | 0.0579 |
|  | Carcass Sex/TRIS * Carcass Infection Status | 2 | 4.498866 | 0.1055 |
| **Larval DCV Titre** | Carcass Sex | 1 | 0.6965869 | 0.4039 |
|  | Carcass Infection Status | 2 | 6.422816 | 0.0403* |
|  | Carcass Sex * Carcass Infection Status | 2 | 0.2180497 | 0.8967 |
| **Adult DCV Titre** | Carcass Infection Status | 2 | 9.6744207 | 0.0079** |
| **Number of Larvae to Pupate** | Carcass Sex | 1 | 13.273693 | 0.0003*** |
|  | Carcass Infection Status | 2 | 0.0745433 | 0.9634 |
|  | Carcass Sex * Carcass Infection Status | 2 | 0.6184649 | 0.7340 |
| **Number of Pupae to Eclose** | Carcass Sex | 1 | 0.0173846 | 0.8951 |
|  | Carcass Infection Status | 2 | 0.1799928 | 0.9139 |
|  | Carcass Sex * Carcass Infection Status | 2 | 0.1485258 | 0.9284 |

### DCV is transmitted to larvae when scavenging on infected carcasses

DCV was detected in larvae collected from plates containing an infected carcass (Figure 3a, Table 1), confirming that scavenging infected carcasses is a viable route of virus transmission. As expected, larvae surrounding DCV-infected carcasses were found to have significantly higher DCV titres when compared to larvae collected from control plates (which contained only uninfected carcasses). However, we also detected DCV infection in larvae surrounding clean carcasses that were housed in a two-choice plate (containing both infected and uninfected carcasses) (Figure 3a), suggesting that some larvae may have moved between food sources in these plates during the assay.

**Figure 3. Fitness consequences of infectious scavenging.**
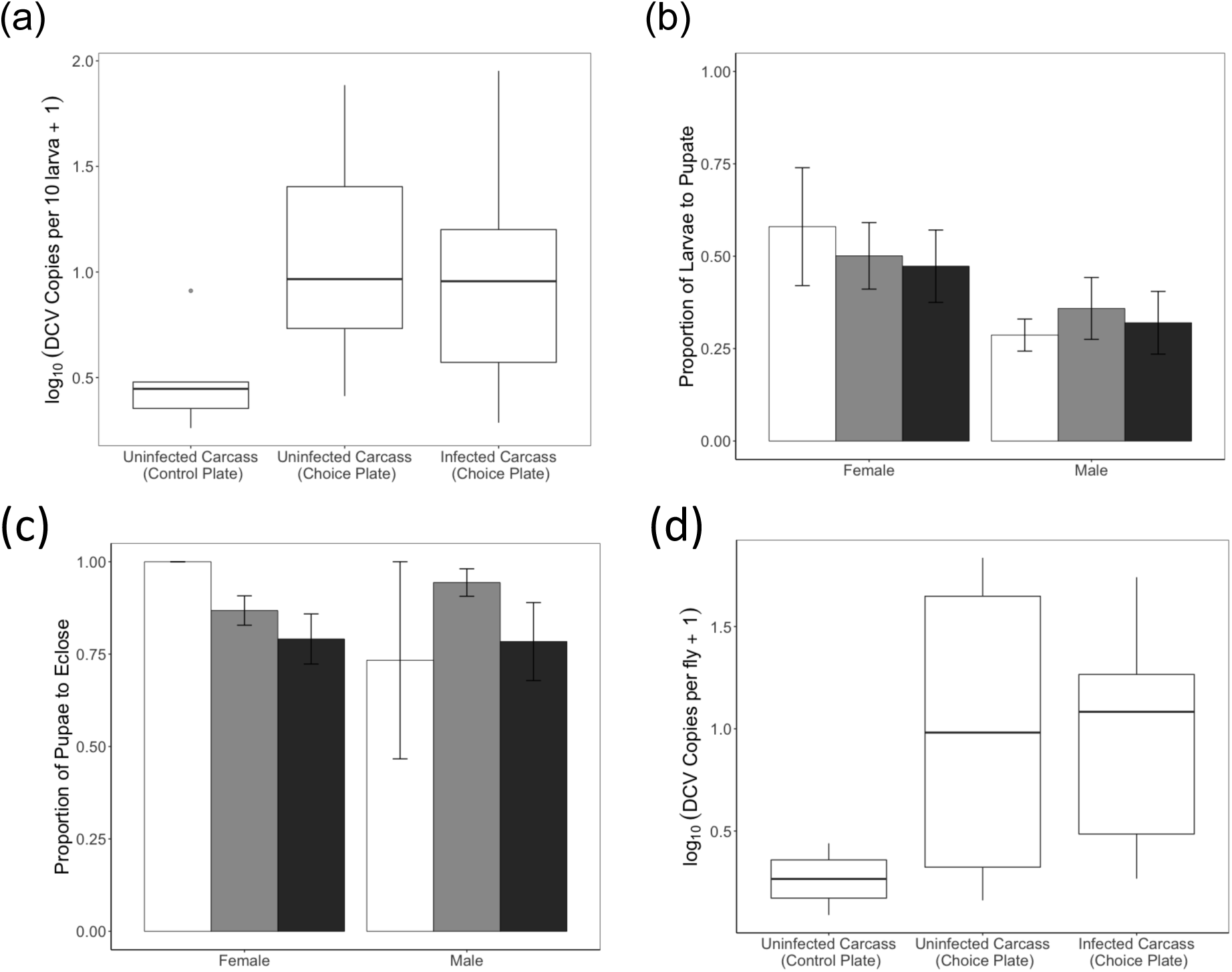
(a) The number of DCV copies present in larvae, quantified immediately after choice assays having fed on an uninfected carcass on a control plate or a choice plate and an infected carcass from a choice plate. Mean ± SE proportion of larvae taken from carcass sites on both choice and control plates to pupate (b) and (c) eclose. Larvae (and the subsequent pupae) were taken from male and female carcasses and varied in their infectious status, an uninfected carcass on a control plate (white bar), an uninfected carcass on a choice plate (grey bar) or an infected carcass on a choice plate (black bar). (d) The number of DCV copies present in adults derived from choice plate assays.

### No effect of virus transmission on larval development

Acquiring infection by scavenging on infectious carcasses had no detectable effect on larval development into pupae (Figure 3b), or in the proportion of pupae that eclosed as adults (Figure 3c; Table 1). However, larval development to pupal stage was significantly higher in larvae that had fed on female carcasses (Figure 3b; Table 1): 50% of larvae feeding on female carcasses reached pupation, while a significantly lower proportion (32%) reached pupation if they had fed on male carcasses (Figure 3b). Following pupation, there was no effect of carcass sex or infection status on the proportion of pupae that eclosed as adults (Figure 3c, Table 1).

### Virus acquired during the larval stage can persist into adulthood

We measured DCV titres in flies that eclosed as adults (Figure 3d). While no DCV infection was detected in flies originally collected near clean carcasses, we detected DCV in 7 out of 11 adult flies that were collected from infected carcasses, suggesting that DCV infection can persist through metamorphosis into the adult insect stage.

### Oviposition preference changes over time and depends on the female’s infection status

Female flies showed a clear preference for oviposition sites containing a carcass, but this choice depended on the fly infection status (Figure 4a, 4b; Table 2). Within the first 24-hour period, uninfected female flies laid significantly more eggs at sites containing a clean carcass compared to sites with an infected carcass or just food (Figure 4a). Female flies infected with DCV, however, did not distinguish between infected and clean carcasses, but still laid significantly fewer eggs at sites without any carcass (Figure 4a). In the 24-48 hour observation period, uninfected females still laid more eggs at sites with carcasses, but no longer preferred the sites containing a clean carcass (Figure 4b; Table 2). DCV-infected females also laid more eggs at sites with an uninfected carcass (pairwise contrast, p<0.0001), but laid even more eggs on sites containing an infected carcass (pairwise contrast, p<0.001) (Figure 4b).

**Figure 4. Adult oviposition choice and fitness consequences.**
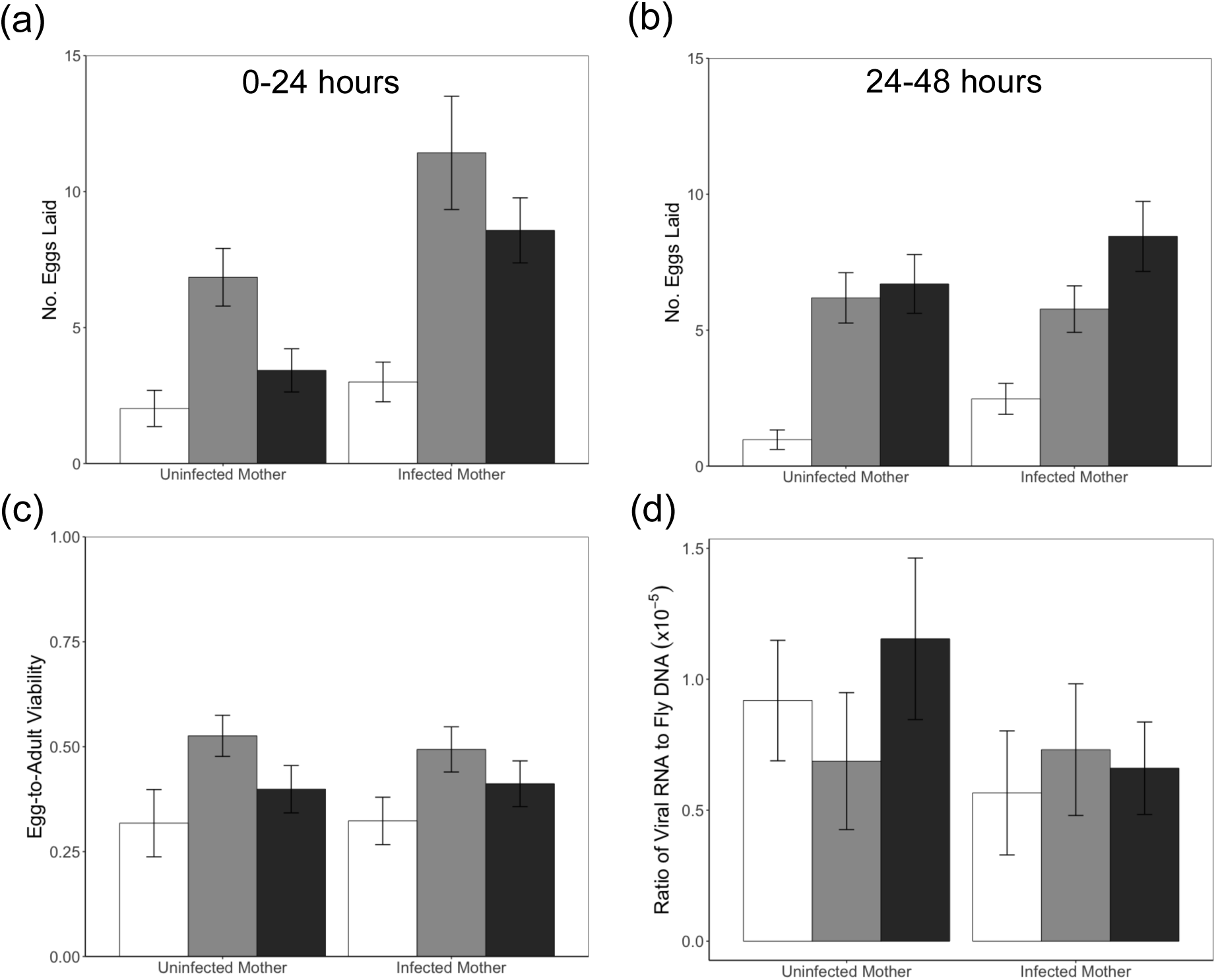
The mean±SE number of eggs laid by infected and uninfected mothers at the three oviposition sites after (a) the first 24 hours of the experiment and (b) the second 24-hour period. (c) The mean ± SE proportion of eggs to develop through to adulthood (egg-to-adult viability) of the clutches laid during the oviposition site choice assay. (d) The mean ± SE ratio of viral RNA to fly DNA in the clutches laid during the oviposition site choice assay. Across all panels, oviposition site treatments are shown using the same colour scheme: food only oviposition sites in white, food and uninfected carcass sites in grey and food and infected fly carcass sites in black.

**Table 2.** Model outputs for statistical tests performed on all experiments testing the causes and costs of infection avoidance in D. *melanogaster* adult oviposition. Significant predictors are marked with asterisks (p<0.05=*, p<0.01=** and p<0.001=***).

| Response Variable | Predictor | DF | F-ratio | p-value |
| --- | --- | --- | --- | --- |
| **Eggs Laid 0-24 hours** | Oviposition Site | 2 | 133.2992 | <0.0001*** |
|  | Maternal Infection Status | 1 | 1.1512 | 0.292 |
|  | Oviposition Site * Maternal Infection Status | 2 | 6.5983 | 0.0042** |
| **Eggs Laid 24-48 hours** | Oviposition Site | 2 | 108.039 | <0.0001*** |
|  | Maternal Infection Status | 1 | 0.00001 | 0.99 |
|  | Oviposition Site * Maternal Infection Status | 2 | 11.278 | 0.0042** |
| **Egg-to-Adult Viability** | Oviposition Site | 2 | 5.6058 | 0.0053** |
|  | Maternal Infection Status | 1 | 0.0128 | 0.88 |
|  | Oviposition Site * Maternal Infection Status | 2 | 0.528 | 0.5917 |
| **Clutch DCV Load** | Oviposition Site | 2 | 2.5523 | 0.0988 |
|  | Maternal Infection Status | 1 | 0.6277 | 0.4359 |
|  | Oviposition Site * Maternal Infection Status | 2 | 1.4596 | 0.2522 |

### Fitness consequences of oviposition preference

Egg-to-Adult viability differed significantly between oviposition sites, and was lower in food-only sites compared to sites containing a carcass (Figure 4c; Table 2). Clutches emerging at carcass sites however, did not differ in their egg-to-adult viability (Figure 4c; Table 2), even though we detected DCV within flies that developed around DCV-infected carcasses (Figure 4d). The infection status of mothers had no effect on egg-to-adult viability (Figure 4c; Table 2) or on the viral load of these clutches (Figure 4d; Table 2).

## Discussion

Viral infection is widespread among invertebrates [32,33], and can cause considerable morbidity and mortality [24,28,34,35]. We should therefore expect selection for mechanisms that allow hosts to detect and avoid infectious conspecifics or potentially infectious environments [3,4]. In the present work, we examined how larval foraging and adult oviposition in *D. melanogaster* are modified in the presence of potential infection by the horizontally transmitted Drosophila C virus (DCV), which is known to cause a variety of physiological and behavioural pathology in fruit flies [24–28].

Our results confirm previous findings that *Drosophila* larvae will actively cannibalise conspecific carcasses when placed in a nutrient-poor environment [17,18], and go further to demonstrate that necrophagy is a viable route for transmission of Drosophila C Virus. The consumption of infectious conspecifics, either through cannibalism or necrophagy, has been demonstrated as a viable route of infection in a wide range of mammalian, amphibian and insect species [36–40]. In holometabolous insects, this phenomenon has been particularly well investigated in Lepidoptera, where cannibalism and/or necrophagy of infected conspecifics has also shown to be a viable route of transmission of several viruses during larval development [39,41–44].

Despite the risk of acquiring infection during cannibalistic foraging, we found no evidence that larval-stage flies could discriminate and avoid infectious carcasses from clean ones. Our findings contrast with a recent study in which *Drosophila* larvae showed evasion of food containing a bacterial suspension of virulent *Pseudomonas entomophila* [45]. Avoidance was no longer observed when using a less virulent strain of the bacterial pathogen, suggesting that external cues about the relative risk and severity of infection are key to avoidance behaviours. The differences in findings likely result from differential olfactory and chemo-sensory factors involved in viral and bacterial detection in Drosophila larvae. Furthermore, while Surendran *et al* (2017) tested evasion in 1^st^ instar larvae, in the current study larval foraging choice was recorded during the 3^rd^ instar, as this is the period of development when foraging activity and feeding is known to peak [46]. Given that larvae are known to actively migrate towards higher quality food [47], the lack of trophic infection avoidance suggests that selection for avoidance of this viral infection is weak. Weak selection for avoidance would be expected if, for example, the fitness costs of DCV infection are low during larval stage infection.

Our data is consistent with a low cost of infection in larvae, as the low titres of DCV acquired during larval feeding on carcasses did not have severe consequences for larval development. Our results contrast with a previous study on DCV infection of larval *D. melanogaster* which reported a 14% reduction in egg-to-adult viability, and severe mortality in adults emerged from infected larvae [26]. Larva in that study were exposed to a highly-concentrated homogenate of DCV-infected flies, and exposed continuously during development until 4-days post-eclosion. This difference in viral exposure may explain the more severe costs of DCV infection compared to this study.

In contrast to the lack of discrimination seen during larval foraging, we found that adult female flies do discriminate between different types of oviposition sites. Uninfected female flies laid more eggs on sites containing an uninfected or infected carcass and food, than a site comprised only of food despite the infection risk this presents. One possible reason for this apparently risky strategy is that while a conspecific carcass can present an infection risk it is also a potential source of additional nutrition [48]. Starved *D. melanogaster* larvae assess the nutritional value of carcasses, ranging from conspecifics to natural predators (Ahmad *et al.*, 2015), and tune their foraging strategies accordingly to optimally forage. Clutches developing on oviposition sites with a carcass present had significantly higher egg-to-adult viability than food only sites (Figure 4c). The preference we see for oviposition sites containing a carcass may therefore indicate that the nutritional value of carcasses on the oviposition sites, rather than infection risk, is driving oviposition-site preference.

During the first 24 hours of egg laying, uninfected flies laid significantly more eggs around uninfected carcasses. This suggests that the presence of DCV is being detected and avoided during oviposition. It is unclear which cues of DCV are detected by females, whether they are detecting the virus directly, or cues of virus derived pathology in the fly carcass. Similar avoidance of pathogenic bacteria has been described in both *D. melanogaster* [6,8,10] and *C. elegans* [49,50]. Avoidance of virus infection has also been described in a range of invertebrates, such as gypsy moth larvae that avoid eating leaves contaminated with virus [51] and lobsters that avoid virus-infected conspecifics [52]. This avoidance likely relies on dedicated chemosensory pathways for olfactory cues [6,9,10,49].

Following the initial 24-hour period, this preference for uninfected carcasses was no longer observed (Figure 4b). We interpret this shift in oviposition-site preference as the result of a trade-off faced by females between minimising DCV infection risk and maximising fecundity. The finite nutritional value of each oviposition site dictates an optimal clutch size that each site can support. If females exceed this, fewer resources are available per offspring. As uninfected flies laid more eggs on non-infectious carcass sites in the first 24 hours, the optimal clutch size is approached sooner than the other two sites. Fruit flies integrate the nutritional quality of oviposition sites into deciding between laying more eggs and acquiring more resources to develop more eggs [53], a trade-off that is also seen in a range of other organisms [48,53–55]. In order to maximise the number of eggs laid, females therefore appear to risk DCV infection by laying their eggs near an infected carcass. The relative nutritional value and the potential costs of DCV infection are patent in the egg-to-adult viability of offspring from each oviposition site: the increase in viability between the food-only site and both the uninfected and infected carcass sites reflects the nutritional difference between these sites. As is clear from Figure 4c, the benefits of oviposition near any carcass appear to outweigh the potential costs of virus infection.

In contrast to uninfected females, females infected with DCV did not discriminate between infectious and non-infectious carcasses, laying the same number of eggs in either oviposition site (Figure 4a,b). Furthermore, in the second 24-hour period, infected females laid significantly more eggs at infectious carcass sites. We interpret this difference in discrimination between infected and healthy females as being driven by the mother’s, rather than the offspring infection risk. For infected females already paying the cost of infection, there is little benefit from avoiding infectious sites.

In summary, our results show that *D. melanogaster* larvae and adults respond to infection risk differently during foraging and oviposition. Notably, oviposition site choice was affected by the female’s infection status and the time-dependent nutritional value of oviposition sites. The initial DCV avoidance shown by mothers during oviposition may also explain why larvae do not avoid DCV during foraging. Alongside a relatively low cost of infection, larvae simply may not need to avoid infection because their mothers have evolved to avoid infectious sites where possible during oviposition. As larvae are not able to forage over large distances, their development - and ultimately their fitness - relies heavily on their mother’s capacity to pick the environment that maximises nutritional value while minimising the risk of infection.

